# Mechanosensitive channel inhibition attenuates TGFβ2-induced actin cytoskeletal remodeling and reactivity in mouse optic nerve head astrocytes

**DOI:** 10.1101/2021.06.05.447194

**Authors:** Alexander Kirschner, Ana N. Strat, John Yablonski, Tyler Bagué, Haiyan Li, Samuel Herberg, Preethi S. Ganapathy

**Affiliations:** Department of Ophthalmology & Visual Sciences, SUNY Upstate Medical University, Syracuse, NY 13210, USA; Department of Neuroscience and Physiology, SUNY Upstate Medical University, Syracuse, NY 13210, USA; Department of Cell and Developmental Biology, SUNY Upstate Medical University, Syracuse, NY 13210, USA; BioInspired Institute, Syracuse University, Syracuse, NY 13244, USA; Department of Biochemistry and Molecular Biology, SUNY Upstate Medical University, Syracuse, NY 13210, USA; Department of Biomedical and Chemical Engineering, Syracuse University, Syracuse, NY 13244, USA

**Keywords:** transforming growth factor beta 2, extracellular matrix, cross-linked actin networks, CLAN, glaucoma, POAG, gliosis, fibronectin

## Abstract

Astrocytes within the optic nerve head undergo actin cytoskeletal rearrangement early in glaucoma, which coincides with astrocyte reactivity and extracellular matrix (ECM) deposition. Elevated transforming growth factor beta 2 (TGFβ2) levels within astrocytes have been described in glaucoma, and TGFβ signaling induces actin cytoskeletal remodeling and ECM deposition in many tissues. A key mechanism by which astrocytes sense and respond to external stimuli is via mechanosensitive ion channels. Here, we tested the hypothesis that inhibition of mechanosensitive channels will attenuate TGFβ2-mediated optic nerve head astrocyte actin cytoskeletal remodeling, reactivity, and ECM deposition. Primary optic nerve head astrocytes were isolated from C57BL/6J mice and cell purity was confirmed by immunostaining. Astrocytes were treated with vehicle control, TGFβ2 (5 ng/ml), GsMTx4 (a mechanosensitive channel inhibitor; 500 nM), or TGFβ2 (5 ng/ml) + GsMTx4 (500 nM) for 48 h. FITC-phalloidin staining was used to assess the formation of f-actin stress fibers and to quantify the presence of crosslinked actin networks (CLANs). Cell reactivity was determined by immunostaining for GFAP. Levels of fibronectin deposition were also quantified. Primary optic nerve head astrocytes were positive for the astrocyte marker GFAP and negative for markers for microglia (Iba1) and oligodendrocytes (OSP1). Significantly increased %CLAN-positive cells were observed after 48-h treatment with TGFβ2 vs. control in a dose-dependent manner. Co-treatment with GsMTx4 significantly decreased %CLAN-positive cells vs. TGFβ2 treatment and the presence of f-actin stress fibers. TGFβ2 treatment significantly increased GFAP and fibronectin fluorescence intensity, which were decreased with GsMTx4 treatment. Our data suggest inhibition of mechanosensitive channel activity as a potential therapeutic strategy to modulate actin cytoskeletal remodeling within the optic nerve head in glaucoma.

## 1. Introduction

Glaucoma is a leading cause of irreversible blindness, characterized by retinal ganglion cell death (Quigley and Broman 2006, Tham, Li et al. 2014). A key site of glaucomatous ganglion cell injury is at the optic nerve head (ONH) (Anderson and Hendrickson 1974, Quigley and Anderson 1976, Quigley, Guy et al. 1979). Although there are differences in ONH structure across species, the matrix and cellular composition are remarkably consistent (Morrison, Jerdan et al. 1988, Howell, Libby et al. 2007, Morrison, Cepurna Ying Guo et al. 2011). This region contains astrocytes, lamina cribrosa cells and microglia in an extracellular matrix (ECM) scaffold that provides structural support to retinal ganglion cell axons as they exit the globe (Hernandez, Luo et al. 1987, Hernandez, Igoe et al. 1988). In glaucoma, the ONH ECM structure undergoes remodeling and stiffening, which correlates with mechanical insult on ganglion cell axons (Crawford Downs, Roberts et al. 2011, Voorhees, Jan et al. 2017, Liu, McNally et al. 2018, Pijanka, Markov et al. 2019, Hopkins, Murphy et al. 2020). However, the mechanism of this stiffening response remains unclear. One likely modifier of ECM structure is the profibrotic cytokine transforming growth factor beta 2 (TGFβ2). Elevated levels of TGFβ2 have been documented in the glaucomatous ONH, and TGFβ2 directly increases ECM deposition by ONH cells (Pena, Taylor et al. 1999, Fuchshofer, Birke et al. 2005, Zode, Sethi et al. 2011).

Due to their close interactions with the ECM and retinal ganglion cell axons, ONH astrocytes are the most likely sensors of pathologic stimuli (Hernandez 2000, May and Lutjen-Drecoll 2002, Morrison, Cepurna Ying Guo et al. 2011). Thus, they are ideally positioned to sense stressors and transduce this insult into changes in ECM makeup and ganglion cell axonal health. In response to glaucomatous injury, astrocytes immediately reorganize their cytoskeletal structure and undergo significant process reorientation (Cooper, Collyer et al. 2018, Tehrani, Davis et al. 2019, Cooper, Pasini et al. 2020). These cytoskeletal changes coincide with the development of astrocyte reactive gliosis, which correlates with cellular hypertrophy and upregulation of glial fibrillary acid protein (GFAP) (Cooper, Collyer et al. 2018). There is clear evidence that astrocyte mobilization and reactive gliosis are protective to retinal ganglion cells in experimental models of glaucoma (Sun, Moore et al. 2017, Cooper, Pasini et al. 2020). However, it is possible that the later effects of unrestricted astrocyte gliosis ultimately contribute to ONH ECM dysregulation and subsequent ganglion cell insult.

A key mechanism by which cells sense and respond to external stimuli is via mechanosensitive ion channels (Petho, Najder et al. 2019). Both Transient Receptor Protein and Piezo channels have been identified in the ONH (Choi, Sun et al. 2015). In response to mechanical stimuli, these channels are activated by direct lipid-stretch or via integrin-mediated stimulation to permit calcium influx to effect downstream signaling (Petho, Najder et al. 2019). In normal conditions, these channels respond to mechanical cues to guide developmental and physiologic cellular organization. By contrast, in pathologic states, dysregulated astrocyte mechanosensitive channel activity alters the actin network, ultimately promoting mechanosensitive channel expression via a positive feedback loop (Chen, Wanggou et al. 2018). Indeed, Piezo1 stimulation directly induces ONH astrocyte gliosis (Liu, Yang et al. 2021); yet, the role of astrocyte mechanosensation in glaucoma is not well understood.

Taken together, TGFβ2 is strongly linked to astrocyte cytoskeletal remodeling and reactive gliosis, and mechanosensitive channel activity can directly regulate actin cytoskeletal reorganization. Thus, modulation of mechanosensitive channels represents an attractive strategy to target astrocyte behavior in glaucoma. In the present study, we tested the hypothesis that inhibition of mechanosensitive channels attenuates TGFβ2-induced actin reorganization and reactive gliosis in mouse ONH astrocytes.

## 2. Methods

### 2.1 Mouse ONH astrocyte isolation and culture

Maintenance and treatment of animals adhered to the institutional guidelines for the humane treatment of animals (IACUC #473) and to the ARVO Statement for the Use of Animals in Ophthalmic and Vision Research. Isolation of primary mouse ONH astrocytes was performed according to protocols modified from Zhao et al (Zhao, Mysona et al. 2017) and Mandal et al (Mandal, Shahidullah et al. 2010). Briefly, C57BL/6J mice were purchased from the Jackson Laboratory (Bar Harbor, ME). Six mice aged 6-8 weeks were used for each isolation. Using a SMZ1270 stereomicroscope (Nikon Instruments, Melville, NY), ONH tissue was dissected from each globe proximal to the sclera, with care to discard as much myelinated optic nerve tissue and peripapillary sclera as possible. ONH samples were digested for 15 minutes using 0.25% trypsin (Invitrogen, Carlsbad, CA) at 37°C and resuspended in ONH astrocyte growth medium (Dulbecco’s modified Eagle’s medium, DMEM/F12 (Invitrogen) + 10% fetal bovine serum (Atlanta Biologicals, Atlanta, GA) + 1% penicillin/streptomycin (Invitrogen) + 1% Glutamax (Invitrogen) + 25 ng/ml epidermal growth factor (Sigma, St. Louis, MO). ONH tissue was then plated in cell culture flasks coated with 0.2% gelatin (Sigma) and maintained at 37°C in a humidified atmosphere with 5% CO_2_. ONH astrocytes were allowed to migrate from the tissue and were passaged after 10-14 days. Three separate cultures were used for the experiments within this study at passages 2 – 5.

### 2.2 ONH astrocyte characterization

ONH astrocytes were seeded at 1 x 10^4^ cells/cm^2^ on sterilized glass coverslips in 24-well culture plates (Thermo Fisher Scientific, Waltham, MA). After 48 h, cells were fixed with 4% paraformaldehyde (PFA; Thermo Fisher Scientific) at room temperature for 10 min, and permeabilized with 0.5% Triton X-100 (Thermo Fisher Scientific) at room temperature for 30 min. Cells were washed in PBS, and blocked (PowerBlock; Biogenx, San Ramon, CA) for 1 h at room temperature. Cells were then incubated for 1 h at room temperature with rabbit anti-glial fibrillary acidic protein (GFAP, 1:300; Dako, Carpinteria, CA), rabbit anti-oligodendrocyte specific protein (OSP, 1:100; Abcam, Cambridge, MA) or rat anti-F4/80 (1:50; BioRad, Hercules, CA). Cells were again washed in PBS, and incubated for 1 h at room temperature with Alexa Fluor^®^ 488-conjugated secondary antibodies (1:500; Abcam). Nuclei were counterstained with 4’,6’-diamidino-2-phenylindole (DAPI; Abcam). Coverslips were mounted with ProLong™ Gold Antifade (Thermo Fisher Scientific) on Superfrost™ Plus microscope slides (Fisher Scientific) and fluorescent images were acquired with an Eclipse N*i* microscope (Nikon). Four representative fields at 20x magnification were taken from each coverslip per culture; number of GFAP-, OSP-, or F4/80-positive cells versus total number of cells were quantified.

### 2.3 Cell treatments

ONH astrocytes were seeded at 1 x 10^4^ cells/cm^2^ on sterilized glass coverslips in 24-well culture plates (Thermo Fisher Scientific, Waltham, MA). After 24 h, cells were treated with increasing doses of TGFβ2 (vehicle control, 1.25 ng/ml, 2.5 ng/ml, and 5 ng/ml; R&D Systems, Minneapolis, MN) for 48 h. In a separate set of experiments, cells underwent the following treatments for 48 h: (1) vehicle control, (2) TGFβ2 (5 ng/ml), (3) TGFβ2 (5 ng/ml) + GsMTx4 (500 nM; Sigma-Aldrich), or (4) GsMTx4 (500 nM).

### 2.4 Staining for f-actin and analysis of cross-linked actin networks (CLANs)

After treatment for 48 h, coverslips were stained for <u>f</u>ilamentous <u>f</u>-actin as previously described (Li, Bague et al. 2021). Briefly, cells were fixed with 4% PFA, permeabilized with 0.5% Triton™ X-100, and stained with Phalloidin-iFluor 488 (Abcam) according to the manufacturer’s instructions. Nuclei were counterstained with DAPI, and fluorescent images were acquired with an Eclipse N*i* microscope (Nikon). Ten representative images were obtained at 60x magnification for each coverslip. CLANs were identified according to established protocols (Hoare, Grierson et al. 2009, Filla, Schwinn et al. 2011). Specifically, CLANs were identified by their geodesic architecture with triangulation between f-actin spokes and hubs; a minimum of 3-5 hubs was necessary for identification of a CLAN. The number of CLAN-positive cells versus total number of cells was quantified.

### 2.5. Staining and fluorescence quantification of GFAP and fibronectin immunoreactivity

Cells were immunostained for GFAP (rabbit anti-GFAP, 1:300, Dako) and fibronectin (rabbit anti-fibronectin antibody, 1:500, Abcam) as detailed in *Methods 2.2*. Fluorescent intensity was quantified in 10 fields at 20x magnification per coverslip with background subtraction using Fiji software (NIH, Bethesda, MD).

### 2.6. Statistical analysis

Individual sample sizes are specified in each figure caption. Comparisons between groups were assessed by one-way analysis of variance (ANOVA) with Tukey’s multiple comparisons *post hoc* tests, as appropriate. All data are shown with mean ± SD, some with individual data points. For CLAN and fluorescence quantification, each data set represents the mean ± SD of 10 fields of view; the grand means and subsequent ANOVA are depicted. The significance level was set at p<0.05 or lower. GraphPad Prism software v9.1 (GraphPad Software, La Jolla, CA, USA) was used for all analyses.

## 3. Results

### 3.1 ONH astrocyte characterization

Primary ONH astrocytes were derived from C57BL/6J mice aged 6-8 weeks using previously described methods (Mandal, Shahidullah et al. 2010, Zhao, Mysona et al. 2017). Cells were ~97% positive for the astrocyte marker GFAP, and effectively negative for markers of oligodendrocytes or microglia (~3-5%) (Fig. 1), thereby identifying them as ONH astrocytes.

**Figure 1.**
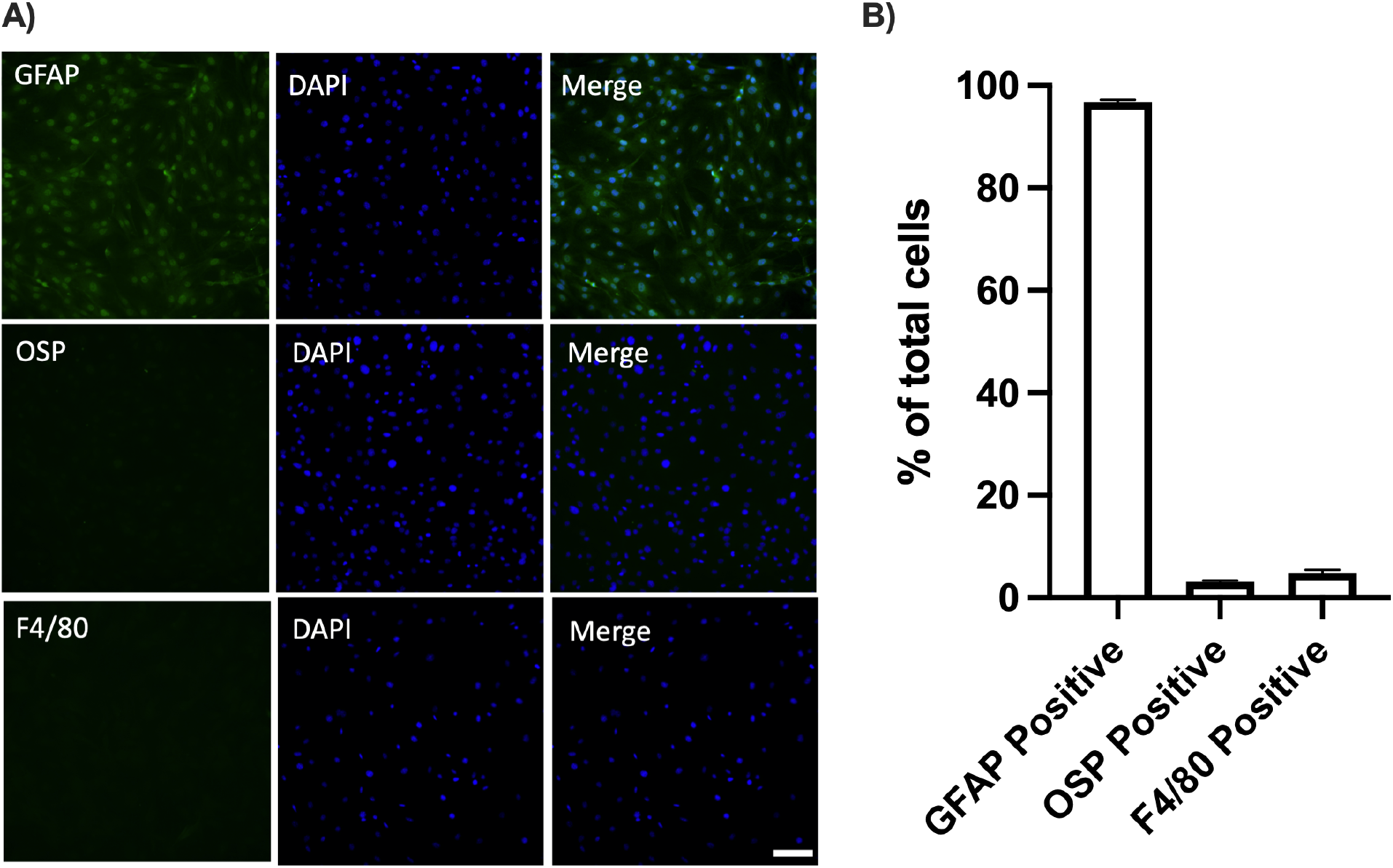
Characterization of primary mouse ONH astrocytes. **(A)** Representative fluorescence images of GFAP (glial fibrillary acid protein), OSP (oligodendrocyte specific protein), and F4/80 (specific for microglia/macrophages) immunostaining. Scale bar 100 μm. (B) Quantification of antibody-positive cells per total number of cells (GFAP 96.8 ± 0.4%; OSP 3.1 ± 0.2%; F4/80 4.8 ± 0.6%). N = 3 groups x 3 biological replicates.

### 3.2 TGFβ2 treatment induces actin cytoskeletal disorganization and CLAN formation in ONH astrocytes

Levels of TGFβ2 are increased within the glaucomatous ONH (Pena, Taylor et al. 1999). Since TGFβ2 induces actin cytoskeletal reorganization in other ocular tissues (Fuchshofer and Tamm 2012, Montecchi-Palmer, Bermudez et al. 2017), we asked whether exogenous treatment of TGFβ2 would exert similar effects on cultured mouse ONH astrocytes.

Vehicle control-treated astrocytes demonstrated baseline organization of the f-actin network (Fig. 2A). Treatment with TGFβ2 altered f-actin fiber morphology (Fig. 2B-D). Specifically, we observed focal areas of disorganized actin fibers, akin to the cross-linked actin networks (CLANs) seen in glaucomatous trabecular meshwork cells (insets Fig. 2C & 2D). To our knowledge, this is the first description of CLAN formation within ONH astrocytes. We found a significant increase in CLAN-containing cells in a dose-dependent manner (Fig. 2E). Treatment with 5 ng/ml TGFβ2 resulted in CLAN formation within ~25% ONH astrocytes; thus 5 ng/ml TGFβ2 was selected for all future treatments.

**Figure 2.**
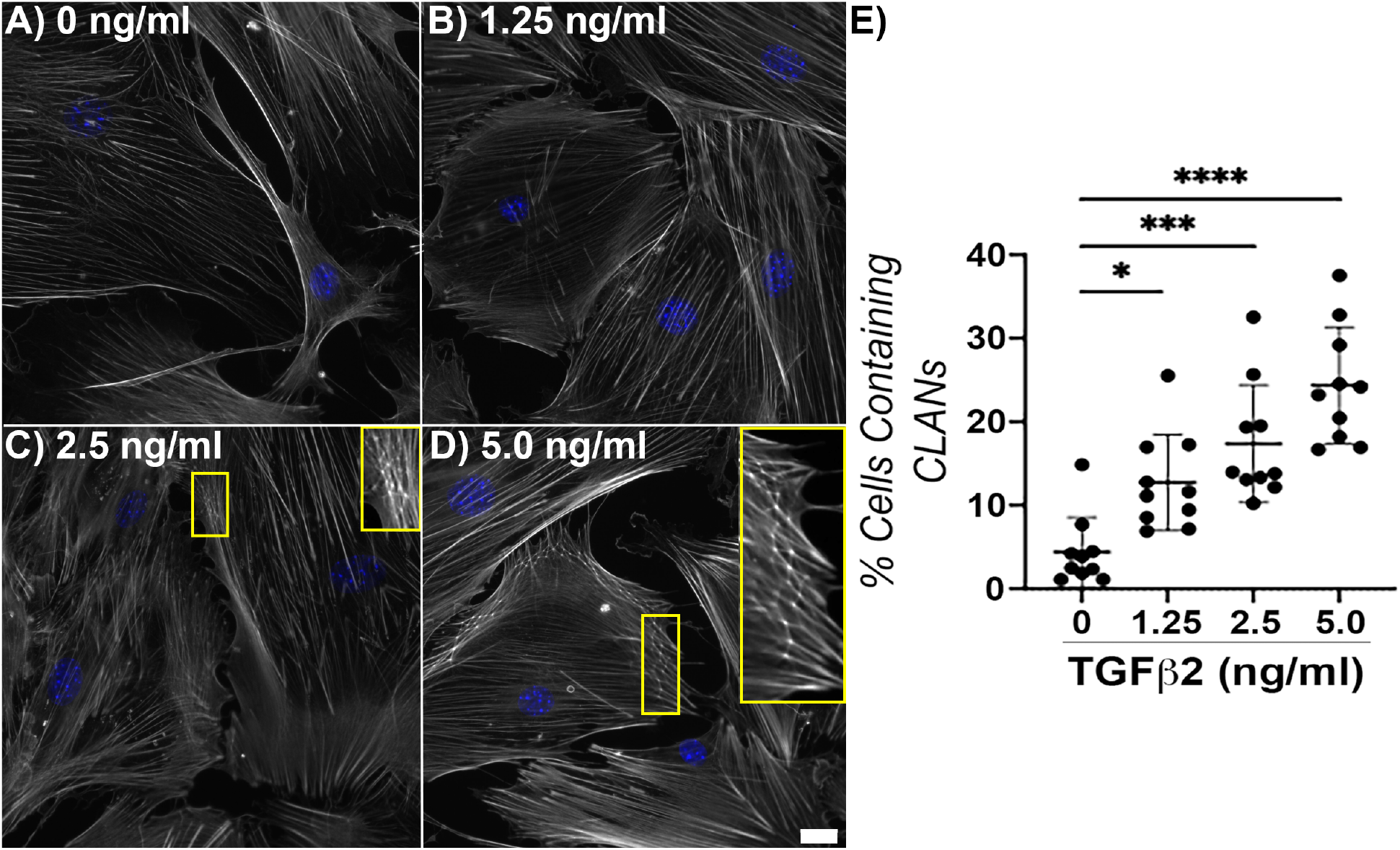
Effect of TGFβ2 treatment on f-actin network. Representative fluorescence images of f-actin within ONH astrocytes following treatment with **(A)** vehicle control, **(B)** TGFβ2 (1.25 ng/ml), **(C)** TGFβ2 (2.5 ng/ml); inset magnified view CLANs, and **(D)** TGFβ2 (5.0 ng/ml); inset CLANs for 48 h (f-actin = white, DAPI = blue). Scale bar 20 μm. **(E)** Quantification of percent cells containing CLANs after TGFβ2 treatment for 48h (vehicle control, 4.38 ± 4.18; 1.25 ng/ml, 12.72 ± 5.78; 2.5 ng/ml, 17.35 ± 7.02; 5.0 ng/ml, 24.35 ± 6.96; * p<0.05; *** p<0.001; **** p<0.0001) N = 3-4 groups x 3 biological replicates.

### 3.3 Co-treatment with a mechanosensitive channel inhibitor reduces TGFβ2-mediated actin cytoskeletal disorganization and CLAN formation in ONH astrocytes

Mechanosensitive channels are critical for cells to sense and respond to external stimuli (Petho, Najder et al. 2019). Therefore, we tested whether co-treatment with the nonspecific mechanosensitive channel inhibitor GsMTx4 (Gnanasambandam, Ghatak et al. 2017) would ameliorate TGFβ2-mediated actin cytoskeletal disorganization in ONH astrocytes.

Again, minimal CLANs were visualized in vehicle control-treated cells (Fig. 3A), while increased f-actin stress fibers and significant CLANs were noted in cells treated with TGFβ2 (Fig. 3B,E). Co-treatment with TGFβ2 and GsMTx4 resulted in f-actin fiber orientation similar to vehicle control-treated cells, and a significantly decreased percentage of CLANs vs. TGFβ2 (12% vs. 25%, p<0.001) (Fig. 3C,E). Lastly, treatment with GsMTx4 also resulted in f-actin fiber morphology comparable to control cells (Fig. 3D,E). These data suggest that mechanosensitive channel inhibition has the potential to modify the ONH astrocyte actin cytoskeleton in response to a glaucomatous stressor.

**Figure 3.**
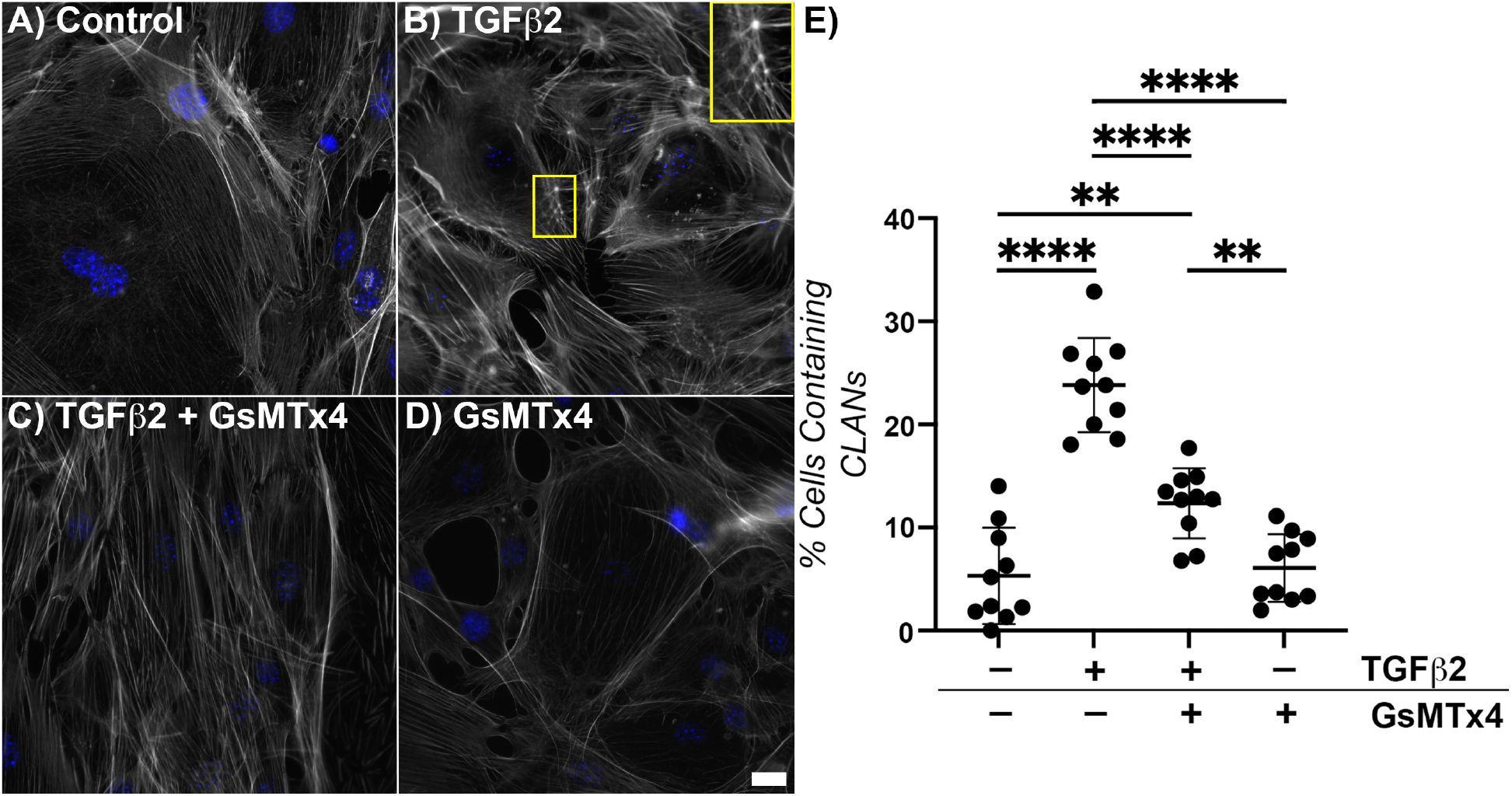
Effect of mechanosensitive channel inhibition on TGFβ2-induced f-actin dysregulation. Representative fluorescence images of f-actin within ONH astrocytes following treatment with **(A)** vehicle control, **(B)** TGFβ2 (5.0 ng/ml); inset CLAN, **(C)** TGFβ2 (5.0 ng/ml) + GsMTx4 (500 nM), and **(D)** GsMTx4 (500 nM) for 48 h (f-actin = white, DAPI = blue). Scale bar 20 μm. **(E)** Quantification of percent cells containing CLANs after above treatments for 48 h (vehicle control, 5.32 ± 4.65; TGFβ2, 23.83 ± 4.56; TGFβ2 + GsMTx4, 12.36 ± 3.39; GsMTx4, 6.07 ± 3.29; ** p<0.01; **** p<0.0001). N = 3-4 groups x 3 biological replicates.

### 3.4 Co-treatment with a mechanosensitive channel inhibitor reduces TGFβ2-mediated GFAP immunoreactivity in ONH astrocytes

Astrocytes within the glaucomatous ONH develop reactive gliosis, associated with changes in their morphology and transcriptional behavior (Sun, Moore et al. 2017). Therefore, we asked whether mechanosensitive channel inhibition would attenuate TGFβ2-stimulated reactive gliosis in ONH astrocytes.

ONH astrocytes treated with vehicle control for 48 h demonstrated low baseline levels of GFAP expression (Fig. 4A,E). Treatment with TGFβ2 induced a significant ~20% increase in GFAP immunofluorescence intensity in cells exposed to TGFβ2 vs. controls (Fig. 4B,E), while co-treatment with GsMTx4 significantly ameliorated the response to TGFβ2, with GFAP intensity returning to baseline levels (Fig. 4C,E). Treatment of ONH astrocytes with GsMTx4 alone resulted in GFAP levels similar to vehicle control (Fig. 4D,E).

**Figure 4.**
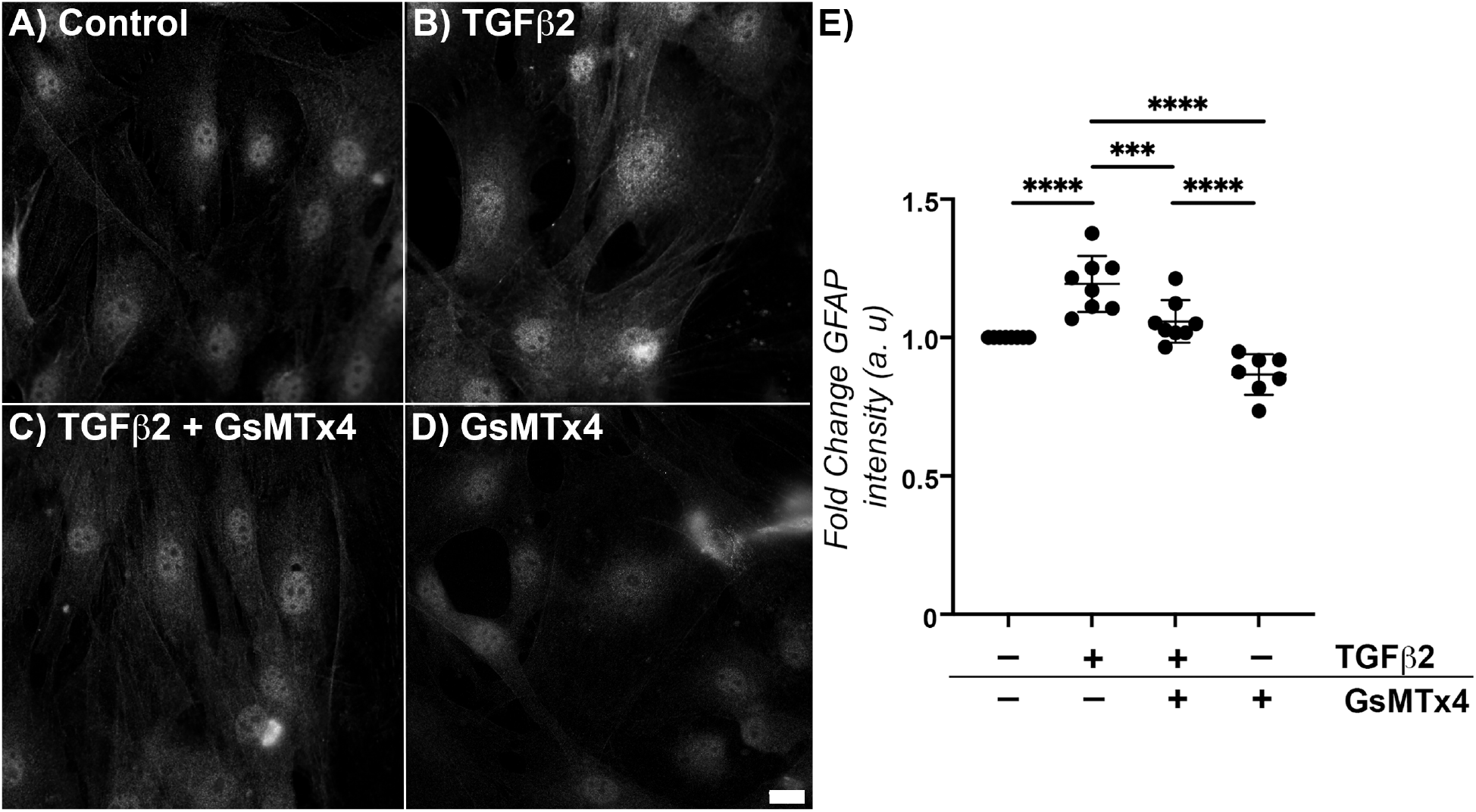
Effect of mechanosensitive channel inhibition on TGFβ2-induced GFAP immunoreactivity. Representative fluorescence images of GFAP within ONH astrocytes following treatment with **(A)** vehicle control, **(B)** TGFβ2 (5.0 ng/ml), **(C)** TGFβ2 (5.0 ng/ml) + GsMTx4 (500 nM), and **(D)** GsMTx4 (500 nM) for 48 h. Scale bar 20 μm. **(E)** Quantification of fold change in fluorescence intensity (grand mean ± SD, arbitrary units) after above treatments for 48 h (vehicle control, 1.00 ± 0.00; TGFβ2, 1.19 ± 0.10; TGFβ2 + GsMTx4, 1.06 ± 0.08; GsMTx4, 0.87 ± 0.07; ** p<0.01; **** p<0.0001). N = 2-3 groups x 3 biological replicates.

### 3.5 Co-treatment with a mechanosensitive channel inhibitor does not alter short-term TGFβ2-mediated fibronectin production by ONH astrocytes

TGFβ2 treatment of ONH astrocytes directly induces production/secretion of fibronectin (Fuchshofer, Birke et al. 2005), which may contribute to overall ECM changes within the ONH. Therefore, we tested whether mechanosensitive channel inhibition would prevent TGFβ2-induced upregulation of fibronectin in ONH astrocytes.

Vehicle control treated ONH astrocytes displayed low levels of fibronectin expression (Fig. 5A,E). Treatment with TGFβ2 for 48 h resulted in a significant ~50% increase in fibronectin intensity vs. controls (Fig. 5B,E). Co-treatment of TGFβ2 and GsMTx4 suggested a decrease in fibronectin levels, but quantitative analyses did not reveal a significant difference (Fig. 5C,E). Treatment with GsMTx4 alone resulted in fibronectin intensity similar to vehicle controls (Fig. 5D,E).

**Figure 5.**
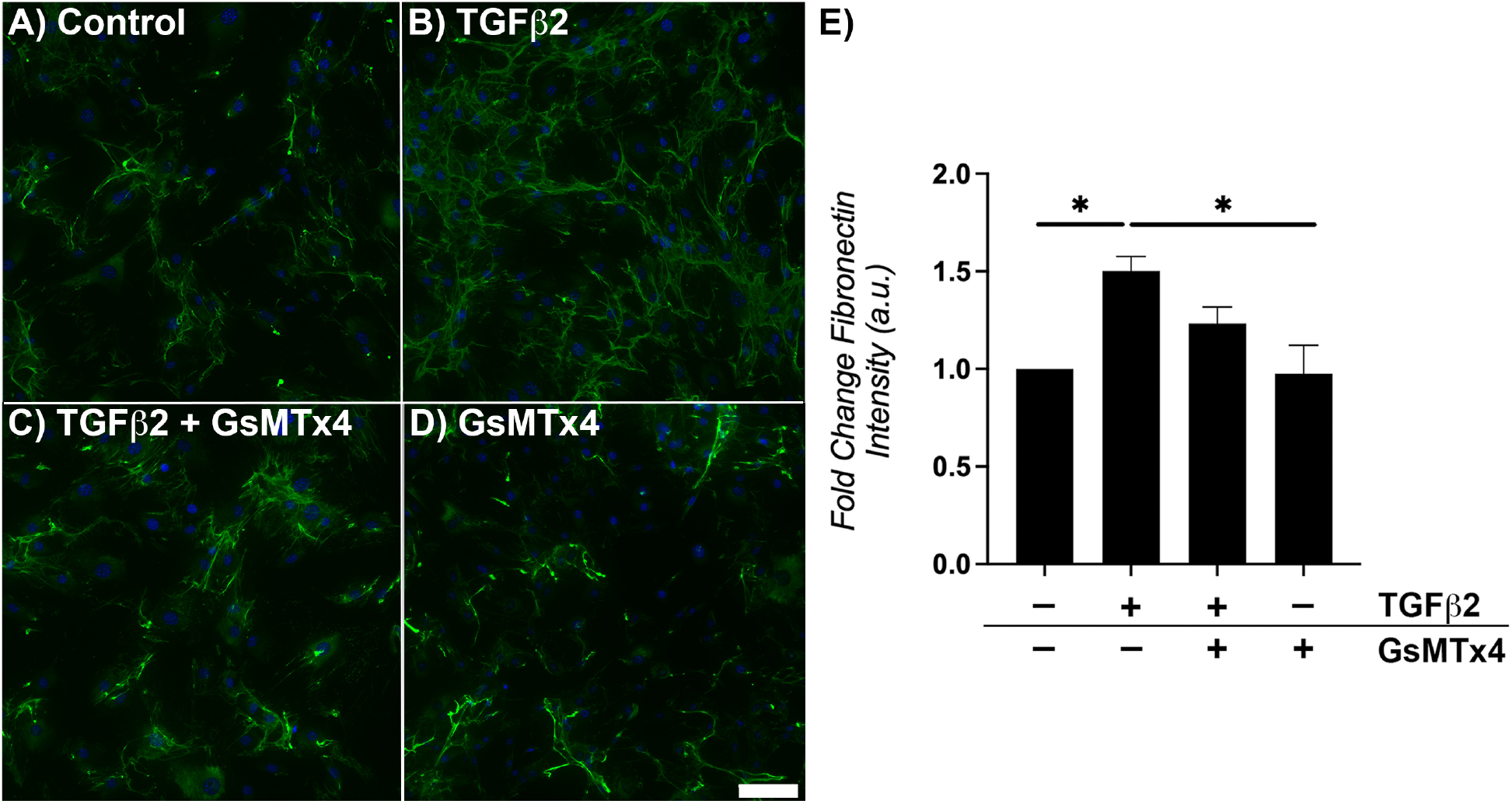
Effect of mechanosensitive channel inhibition on TGFβ2-induced fibronectin immunoreactivity. Representative fluorescence images of fibronectin within ONH astrocytes following treatment with **(A)** vehicle control, **(B)** TGFβ2 (5.0 ng/ml), **(C)** TGFβ2 (5.0 ng/ml) + GsMTx4 (500 nM), and **(D)** GsMTx4 (500 nM) for 48 h (fibronectin = green, DAPI = blue. Scale bar 100 μm. **(E)** Quantification of fold change in fluorescence intensity (grand mean ± SD, arbitrary units) after above treatments for 48 h (vehicle control, 1.00 ± 0.00; TGFβ2, 1.50 ± 0.07; TGFβ2 + GsMTx4, 1.23 ± 0.08; GsMTx4, 0.98 ± 0.15; * p<0.05). N = 3 groups x 2 biological replicates.

## 4. Discussion

Glaucoma is a multifactorial disease, and several mechanisms contribute to eventual retinal ganglion cell dysfunction/loss. Astrocytes have been identified as key responders to glaucomatous insult (Hernandez 2000, Sun, Moore et al. 2017, Cooper, Pasini et al. 2020). In neural tissues, astrocytes modulate synapse formation and mediate metabolic stressors; they are also capable of directing scar formation and remodeling the surrounding ECM (Blanco-Suarez, Caldwell et al. 2017). Within the ONH, astrocyte mobility and reactivity has been shown to promote ganglion cell viability (Sun, Moore et al. 2017, Cooper, Pasini et al. 2020). In certain cases, however, astrocytes may adopt a more detrimental phenotype, becoming neurotoxic (Liddelow, Guttenplan et al. 2017, Sterling, Adetunji et al. 2020), and later in the disease, ONH astrocytes elicit fibrosis of the surrounding ECM, which negatively affects ganglion cell health (Schneider and Fuchshofer 2016). Our overall aim, then, is to elucidate mechanisms governing the complex ONH astrocyte response to glaucomatous stressors, with the ultimate goal of precisely directing ONH astrocyte behavior to best preserve ganglion cell function and viability.

Aging and elevated intraocular pressure are the highest correlated risk factors for glaucoma (Weinreb, Aung et al. 2014). Age-associated stiffening of the outflow pathway within the eye may induce elevated intraocular pressure in susceptible individuals, and this stiffening is linked to increased TGFβ2 levels in that region (Agarwal, Daher et al. 2015). Similarly, TGFβ2 levels are elevated within ONH astrocytes in glaucoma, which may play a role in ONH fibrosis seen in end-stage disease (Pena, Taylor et al. 1999). What is not known, however, is whether TGFβ2 induces astrocyte actin cytoskeletal changes that are seen early in glaucoma, and what mechanisms govern these processes. In the current study, we demonstrate that TGFβ2 treatment of ONH astrocytes elicits alterations to the actin cytoskeleton, and reactive gliosis in a mechanosensitive channel-dependent manner.

Treatment with TGFβ2 resulted in increased actin stress fibers in ONH astrocytes, similar to what is observed in trabecular meshwork cells within the outflow system of the eye (O’Reilly, Pollock et al. 2011), demonstrating the clear potential of TGFβ2 to modulate the ONH astrocyte actin cytoskeleton. Interestingly, we also visualized CLANs (Fig. 2), unique actin arrangements, within ONH astrocytes after treatment. In a single targeted analysis of human optic nerve head tissue, CLANs were not visualized within control nor glaucomatous ONH astrocytes (Job, Raja et al. 2010). It is important to note that cell culture conditions are inherently distinct from the *in vivo* environment, particularly with regard to substrate stiffness and ECM makeup. The mouse ONH astrocytes for these experiments were cultured on uncoated, stiff (megapascal, MPa – gigapascal GPa range) glass substrates. It is possible that these artificial culture conditions simply enhance a naturally more subtle susceptibility when astrocytes reside within their native soft (pascal, Pa – kilopascal, kPa range) ECM. To that end, we recently reported on a 3D hydrogel system to study trabecular meshwork cell behavior in response to glaucomatous stressors (Li, Bague et al. 2021). Future studies will use a version of this ECM biopolymer hydrogel to determine if ONH astrocytes behave similarly in a softer, more biomimetic matrix, and how these actin cytoskeletal changes translate into alterations in astrocyte morphology/mobility. Our data confirm that TGFβ2 induces ONH astrocyte reactivity (Fig. 4) and fibronectin production (Fig. 5); a 3D system will be ideal for investigating how modification in the cell-ECM interface subsequently modulates astrocyte behavior.

The studies described herein are among the earliest to highlight the role of mechanosensitive channels in modulating the ONH astrocyte response to glaucomatous stressors. Previous studies have shown that intraocular pressure elevation in murine models of glaucoma upregulates mechanosensitive channel expression (Choi, Sun et al. 2015), and that mechanical stretch increases Piezo1 activity in ONH astrocytes (Liu, Yang et al. 2021). Here, we showed that co-treatment with the mechanosensitive channel inhibitor GsMTx4 potently reduced TGFβ2-induced actin cytoskeletal dysregulation and CLAN formation (Fig. 3) within a short time-frame (48 h) after exposure to TGFβ2. In addition, GsMTx4 co-treatment reversed TGFβ2-induced reactive gliosis (Fig. 4), which likely modulates retinal ganglion cell viability in glaucoma. GsMTx4 is a relatively nonselective inhibitor of cationic mechanosensitive channels, which include calcium, sodium, and potassium channels (Gnanasambandam, Ghatak et al. 2017). Of these, mechanosensitive calcium channels are attractive candidate mediators of actin fiber realignment since calcium can directly modulate the actin cytoskeleton (Hepler 2016), and future experiments are focused on identifying which key channels are involved.

In conclusion, this study identifies inhibition of mechanosensitive channels as an attractive means to modulate ONH astrocyte behavior in glaucoma. Our data support that mechanosensitive channel inhibition ameliorates features of glaucomatous astrocyte response (i.e., actin cytoskeletal dysregulation and reactive gliosis). As astrocytes are a primary support cell to retinal ganglion cells, it is not yet known how these changes in astrocyte behavior will ultimately affect ganglion cell function. Additional work in our laboratory will explore the impact of astrocyte mechanosensitive channel function on retinal ganglion cell health and function.

## Funding

This project was supported in part by unrestricted grants to SUNY Upstate Medical University Department of Ophthalmology & Visual Sciences from Research to Prevent Blindness (RPB) and from Lions Region 20-Y1, American Glaucoma Society (AGS) Young Clinician Scientist Award to P.S.G. and RPB Career Development Awards to S.H. and P.S.G..

## Acknowledgments

We thank Jing Zhao and Dr. Kathryn Bollinger at Augusta University for guidance in setting up mouse ONH astrocyte culture. We thank Drs. Audrey M. Bernstein and Mariano S. Viapiano, for imaging support. We also thank Dr. Dale D. Hunter for editing this manuscript and Dr. William J. Brunken for his guidance and mentorship.

## Author contributions

A.K., A.N.S., J.Y., T.B., H.L., S.H., and P.S.G designed all experiments, collected, analyzed, and interpreted the data. A.K., A.N.S., and P.S.G. wrote the manuscript. All authors collected data and commented on and approved the final manuscript. S.H. and P.S.G. conceived and supervised the research.

## Competing interests

The authors declare no conflict of interest.

## Data and materials availability

All data needed to evaluate the conclusions in the paper are present in the paper. Additional data related to this paper may be requested from the authors.

## Notes

### Competing Interest Statement

The authors have declared no competing interest.

